# A novel sodium signaling complex regulates uterine activity

**DOI:** 10.1101/2020.07.31.229138

**Authors:** Juan J. Ferreira, Chinwendu Amazu, Lis C. Puga-Molina, Xiaofeng Ma, Sarah K. England, Celia M. Santi

**Author notes:** Both authors are corresponding authors. Correspondence should be addressed to and. Both authors contributed equally to this work (Juan J. Ferreira, Chinwendu Amazu).

## Abstract

Depolarization of the myometrial smooth muscle cell (MSMC) resting membrane potential is necessary for the transition of the uterus from a quiescent state to a contractile state. The molecular mechanisms involved in this transition are not completely understood. Here, we report a novel coupled system between the Na^+^-activated K^+^ channel (SLO2.1) and the non-selective Na^+^ leak channel (NALCN) which determines the MSMC membrane potential. We show that SLO2.1 currents are activated by an inward Na^+^ leak current carried by the NALCN channel leading to MSMC hyperpolarization. These results show an unanticipated role for the Na^+^ leak currents in activating a negative feedback system countering the excitable effects of Na^+^ currents. This is a novel role for the NALCN channel in which Na^+^ acts as an intracellular signaling molecule. In fact, we report here that the net effect of Na^+^ entry through NALCN channels is a hyperpolarization of the MSMCs plasma membrane because of the activation of SLO2.1 K channel. Importantly, we also report that a decrease in NALCN/SLO2.1 activity triggers both Ca^2+^ entries through VDCCs, promoting myometrial contraction. Consistently, with a functional coupling, our data show that NALCN and SLO2.1 are in proximity to one another in human MSMCs. We propose that the spatial arrangement of SLO2.1 and NALCN permits these channels to functionally interact in order to regulate human MSMC membrane potential and cell excitability to modulate uterine contractile activity.

## Introduction

As pregnancy progresses, the uterus transitions from a quiescent state to a highly contractile state at labor. This transition is, in part, driven by changes in the electrical activity of the myometrial smooth muscle cells (MSMCs). At the end of the second trimester, MSMCs have a hyperpolarized (more negative inside the membrane than outside) resting membrane potential of -75 mV, and the membrane potential depolarizes to -50 mV by the time of labor (Parkington, Tonta, Brennecke, & Coleman, 1999). This change of membrane potential is coupled with an increase in the frequency of uterine contractions (Casteels & Kuriyama, 1965; Parkington et al., 1999). The membrane potential is determined primarily by the balance between an outward potassium (K^+^) leak current and an inward sodium (Na^+^) leak current across the plasma membrane. If K^+^ efflux increases, the membrane potential becomes more negative, promoting quiescence. Conversely, it has long been thought that increased Na^+^ influx causes the membrane potential to depolarize, promoting a more excitable and contractile uterine state due to the opening of voltage-gated Ca^2+^ channels. Thus, identifying the ion channels and the molecular mechanisms that control membrane potential is key to understanding how MSMC excitability and uterine contractions are regulated.

We previously reported that SLO2.1, a member of the SLO2 family of Na^+^-activated K^+^ channels, is expressed in human MSMCs (Dryer, 2003; Ferreira et al., 2019; Hage & Salkoff, 2012; Yuan et al., 2003). This channel has low voltage dependence and high conductance and can be significantly active at physiological intracellular Na^+^ concentration (Budelli et al., 2009; Ferreira et al., 2019; Hage & Salkoff, 2012). These characteristics suggest that SLO2.1 can modulate membrane potential and cell excitability by regulating K^+^ efflux. SLO2 channels are also highly expressed in other smooth muscle cells and the brain (Dryer, 2003; Kameyama et al., 1984; P. Li et al., 2019; Smith et al., 2018), where they form complexes with voltage-gated Na^+^ channels (Hage & Salkoff, 2012; Takahashi & Yoshino, 2015). In these complexes, the Na^+^ conducted through the Na^+^ channels modulates K^+^ currents and their effect on membrane potential. Whether SLO2.1 forms a functional complex with a Na^+^ channel in human MSMCs to modulate membrane potential and cell excitability is unknown.

In human MSMCs, the role of voltage-gated Na^+^ channels is unclear, while the voltage-independent Na^+^ leak channel non-selective (NALCN) is expressed in human MSMCs (Reinl, Cabeza, Gregory, Cahill, & England, 2015). NALCN conducts about 50% of the Na^+^ leak current at the membrane potential in human MSMCs (Reinl et al., 2015). Thus, NALCN could be the channel that allows influx of Na^+^ to modulate SLO2.1 and thereby helps regulate membrane potential in MSMCs. Here, we provide evidence that SLO2.1 and NALCN form a novel functional complex that can regulate human MSMC membrane potential, excitability, and uterine contractility.

## Results

### Na^+^ influx through NALCN channels activates SLO2.1 in human MSMCs

To test the hypothesis that Na^+^ influx through NALCN activates SLO2.1, we first performed whole-cell patch-clamp on primary human MSMCs treated with the SLO1 K^+^ channel blocker tetraethylammonium (TEA) (**Figure 1A**) (Khan, Smith, Morrison, & Ashford, 1993). Consistent with our previous findings in MSMCs (Ferreira et al., 2019) and others’ findings in neurons (Hage & Salkoff, 2012; Takahashi & Yoshino, 2015), the addition of 80 mM extracellular Na^+^ led to an 80-100% increase in K^+^ currents at +80 mV and at -60 mV (**Figure 1B**), confirming that SLO2.1 is a Na^+^-activated K^+^ channel. Next, we performed similar experiments in the presence of the Na^+^ leak channel blocker gadolinium (Gd^3+^) and found that 10 µM Gd^3+^ completely inhibited the K^+^ current activated by the addition of extracellular Na^+^ (**Figure 1C-E)**. These results indicate that the Na^+^ that activates K^+^ efflux through SLO2.1 enters MSMCs through a Na^+^ leak channel. To confirm if NALCN channels are the predominant carrier of the Na^+^ that activates SLO2.1, we performed similar experiments using the specific NALCN inhibitor, CP96345 (CP). CP96345 inhibited about 80% of the induced NALCN currents in HEK293 cells (Hahn, Kim, Um, Kim, & Park, 2020). In our hands, CP96345 inhibited over 50% of the endogenous Na^+^ leak current at -60 mV in hTERT cells (**Supp. Figure 2**). The K^+^ current activated by the addition of extracellular Na^+^ was almost completely inhibited by the presence of the NALCN blocker CP96345 (50 µM) in the extracellular solution (**Figure 1H and J)**. Conversely, Gd^3+^ and CP96345 did not affect the SLO2.1 K^+^ current when the intracellular solution contained 80 mM Na^+^ (**Supp. Figure 3)**.

**Figure 1.**
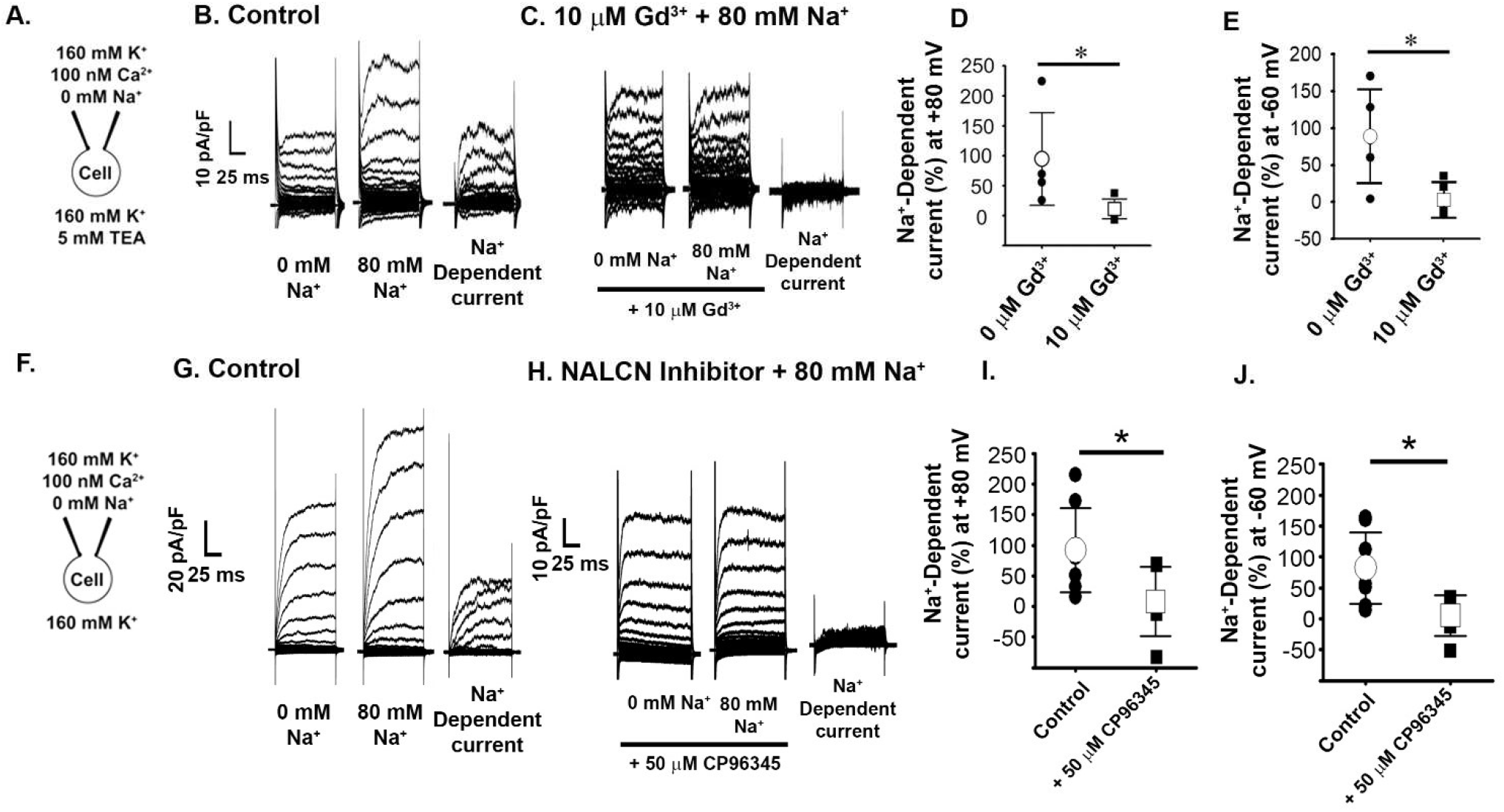
SLO2.1 channels are activated by a NALCN-Dependent Na^+^ leak current in human MSMCs. **(A)** Schematic of whole-cell recording set-up. TEA, tetraethylammonium. **(B)** Representative whole-cell currents (V_h_ = 0 mV, with step pulses from −80 to +150 mV) from human MSMC recorded in 0 mM Na^+^ or 80 mM external Na^+^. The Na^+^-dependent currents were calculated by subtracting traces (80 mM – 0 mM Na^+^). **(C)** Same as **(B)**, in the presence of 10 μM Gd^3+^. **(D** and **E)** Graphs depicting the percentage of Na^+^-dependent currents at +80 mV and -60 mV, respectively, in 0 µM Gd^3+^ or 10 µM Gd^3+^(n = 5 cells, data plotted as the mean and standard deviation). In **(D)**, values are 94.75 ± 77.14 for 0 µM Gd^3+^and 11.20 ± 16.36 for 10 µM Gd^3+^. In **(E)**, values are 89.24 ± 63.56 for 0 µM Gd^3+^and 3.51 ± 23.50 for 10 µM Gd^3+^. **(F)** Schematic of whole-cell recording set-up. **(G)** Representative whole-cell currents (V_h_ = 0 mV, with step pulses from −80 to +150 mV) from hTERT cells recorded in 0 mM Na^+^ or 80 mM external Na^+^. The Na^+^-dependent currents were calculated by subtracting traces (80 mM – 0 mM Na^+^). **(H)** Same as **(G)**, in the presence of 50 μM CP96345 (CP) (NALCN inhibitor). **(I** and **J)** Graphs depicting the percentage of Na^+^-dependent currents at +80 mV and -60 mV, respectively, in 0 µM CP or 60 µM CP. Number of cells is 6 and 8 for 0 and 50 μM CP respectively, data plotted as the mean and standard deviation. In **(I)**, values are 83.09 ± 20.57 for 0 µM CP and 6.50 ± 13.48 for 50 µM CP. In **(J)**, values are 91.59 ± 68.47 for 0 µM CP and 8.19 ± 56.77 for 50 µM CP. * *P*<0.05 by unpaired t-test.

### SLO2.1-induced membrane hyperpolarization is modulated by a NALCN-dependent Na^+^ leak current

Efflux of K^+^ hyperpolarizes the MSMC membrane potential, so we determined whether Na^+^ influx could activate SLO2.1 and thereby lead to membrane hyperpolarization. To measure changes in membrane potential, we performed flow cytometry of MSMCs in the presence of the fluorescent dye DiSC3(5), a cationic voltage-sensitive dye that accumulates on hyperpolarized membranes (Molina et al., 2019; Plasek & Hrouda, 1991; Santi et al., 2010). Treating the cells with 80 mM extracellular Na^+^ led to a 30.48 ± 20.62% increase in DiSC3(5) fluorescence, indicating that membrane hyperpolarization was activated by Na^+^ influx. In contrast, treating the cells with 80 mM choline, an impermeable cation, only led to a 9.91 ± 9.76% increase in DiSC3(5) fluorescence, indicating that the effect of Na^+^ was not due to a change in osmolarity (**Figure 2)**. Additionally, treating the cells with 80 mM lithium (Li^+^), a permeable cation that does not activate SLO2.1, only led to a 2.01 ± 6.54% increase in DiSC3(5) fluorescence (**Figure 2A**), indicating that Na^+^ influx, and not simply a change in membrane potential, activates SLO2.1.

To determine whether NALCN was responsible for the influx of Na^+^, we first treated cells with Gd^3+^. In this condition, the addition of Na^+^ only increased DiSC3(5) fluorescence by 6.01 ± 7.16% (**Figure 2**). This response was similar to the increased DiSC3(5) fluorescence measured in cells treated with choline or Li^+^ (*P* =0.163 and 0.662, respectively) (**Figure 2**). In a separate experiment, we found that, in the presence of Gd^3+^, Na^+^ and choline increased DiSC3(5) fluorescence by similar amounts (5.63 ± 4.57% vs. 6.01± 7.16%, *P* = 0.833) (**Supp Figure 4A**).

**Figure 2.**
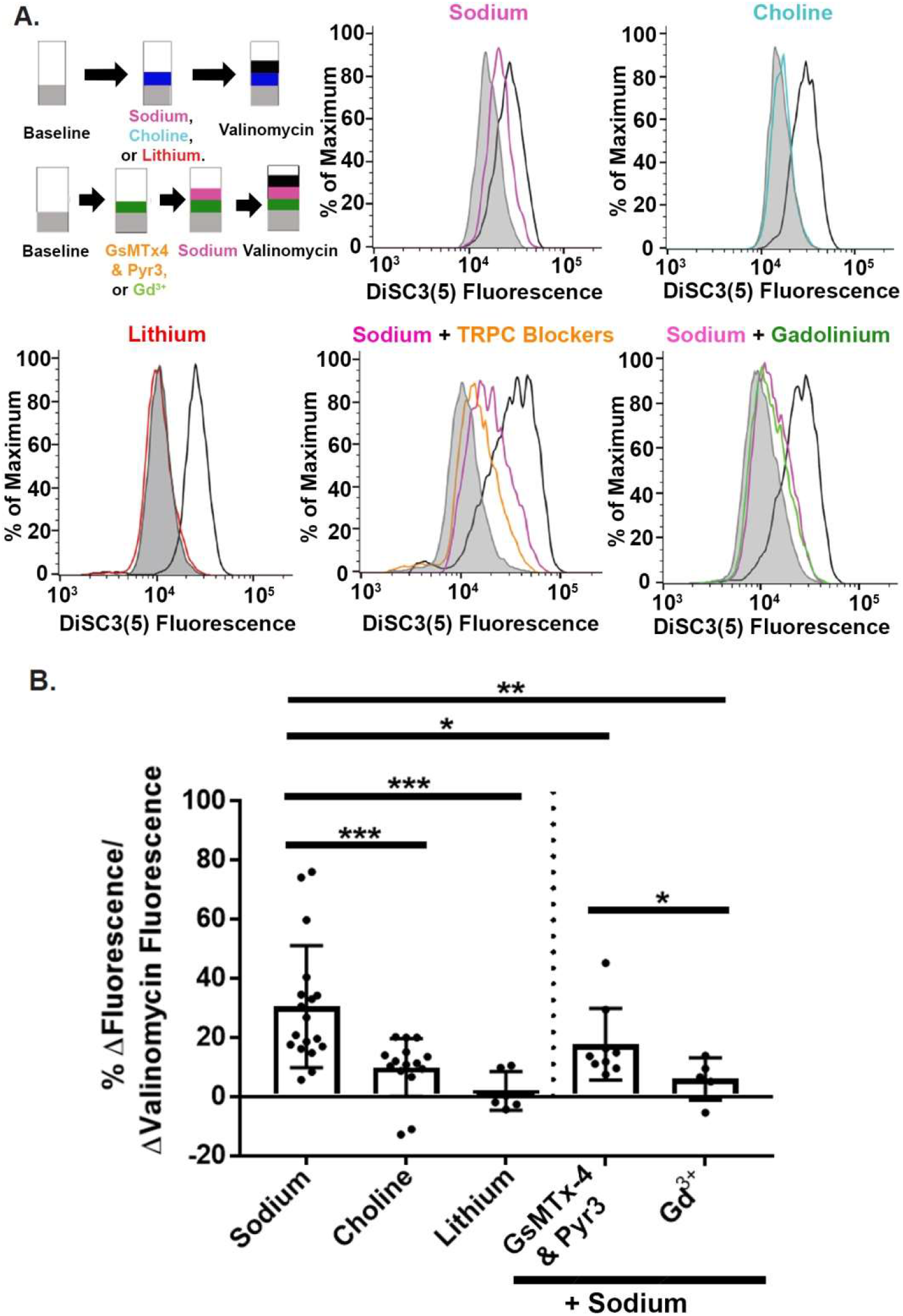
Activation of SLO2.1 by the NALCN-dependent Na^+^ leak hyperpolarizes the membrane potential. **(A)** Experimental schemes and representative images of relative shifts of DiSC3(5) fluorescence induced by sodium, choline, lithium, GsMTx-4, and Pyr3, and Gd^3+^ in hTERT-HM cells. **(B)** Quantification of shifts in cells by sodium (n= 18), choline (n=15), lithium (n=6), and sodium in the presence of GsMTx-4 and Pyr3 (n=9) or Gd^3+^ (n=5) normalized to changes in fluorescence in the presence of valinomycin. Data are presented as mean and standard deviation. \**P*<0.05, \*\**P*<0.01, \*\*\**P*<0.001 by unpaired t-test with Mann-Whitney corrections.

Although Gd^3+^ inhibits NALCN, it also inhibits transient receptor potential canonical (TRPC) channels, which primarily conduct Ca^2+^ but can also conduct Na^+^ (Babich et al., 2004; Dalrymple, Mahn, Poston, Songu-Mize, & Tribe, 2007; Dalrymple, Slater, Beech, Poston, & Tribe, 2002; Ku et al., 2006; Wang et al., 2020). Because TRPC1, 3, and 6 are expressed in human MSMCs (Babich et al., 2004; Dalrymple et al., 2007; Dalrymple et al., 2002; Ku et al., 2006; Wang et al., 2020), we next asked whether GsMTx-4, which inhibits TRPC1 and TRPC6 and Pyr3, which inhibits TRPC3, would affect Na^+^-induced membrane hyperpolarization. In the presence of these inhibitors, Na^+^-induced increase in DiSC3(5) fluorescence was significantly lower than in their absence (17.77 ± 12.10% vs.30.48 ± 20.62%, *P*= 0.040) (**Figure 2**). However, Na^+^-induced increase in DiSC3(5) fluorescence was significantly greater in the presence of these inhibitors than in the presence of Gd^3+^(17.77 ± 12.10% vs. 6.01 ± 7.16%, *P* =0.029) (**Figure 2**). Additionally, in the presence of TRPC blockers, Na^+^ induced a greater increase in DiSC3(5) fluorescence than did choline (17.77 ± 12.10% vs. 5.21 ± 6.426, *P* = 0.008) (**Supp. Figure 4B**). Thus, we conclude that, although Na^+^ influx through TRPC channels is partially responsible for SLO2.1-dependent hyperpolarization, Na^+^ influx through NALCN is responsible for ∼60% of the Na^+^-activated, SLO2.1-dependent hyperpolarization in human MSMCs.

### Functional coupling of Na^+^ influx and K^+^ efflux modulates MSMC Ca^2+^ responses and myometrial contractility

Our data thus far suggested that Na^+^ influx through NALCN activates SLO2.1, leading to K^+^ influx and MSMC membrane hyperpolarization. To determine whether these effects on ion channel activity and membrane potential had a functional outcome, we first examined the effects on intracellular calcium (Ca^2+^). In human MSMCs, membrane depolarization increases Ca^2+^ influx through voltage-dependent Ca^2+^ channels (VDCCs), and Ferreira *et al*. showed that inhibiting SLO2.1 triggered Ca^2+^ entry through VDCCs (Ferreira et al., 2019). Thus, we hypothesized that Na^+^ influx through NALCN activates SLO2.1, leading to K^+^ efflux, membrane hyperpolarization, and inhibition of Ca^2+^ influx.

To test this idea, we used the Ca^2+^ indicator Fluo4-AM to measure intracellular Ca^2+^ concentration. MSMCs at baseline have low-frequency spontaneous calcium release (Lynn, Morgan, Gillespie, & Greenwell, 1993). When we depolarize MSMCs with high concentrations of external KCl (20 and 50 mM), a significant increase in intracellular Ca^2+^ oscillations was observed (**Figure 3A**). Next, to evaluate the effects of extracellular Na^+^ on intracellular Ca^2+^ responses, we treated MSMCs with either 0 mM NaCl plus 160 mM choline (to prevent Na^+^ influx but maintain osmolarity) or 80 mM NaCl plus 80 mM choline (to promote Na^+^ influx). Our results showed that Ca^2+^ oscillations, similar to those induced by 50 mM KCl, were evident in the presence of 0 mM NaCl but not in the presence of 80 mM NaCl (**Figure 3B, C, Supp. Figure 5A)**. These results are in agreement with data reported by Morgan et al,.1993 (Morgan, Lynn, Gillespie, & Greenwell, 1993). We then performed experiments in which we substituted Na^+^ with Li^+^, which does not activate SLO2.1. Ca^2+^ oscillations similar to those induced by 50 mM KCl, were evident in the presence of 135 mM Li^+^ but not in the presence of 135 mM Na^+^ (**Figure 3C and D**). To confirm that the Ca^2+^ increases in 0 mM NaCl (135 mM Li^+^) were dependent on extracellular free Ca^2+^, we repeated the Li^+^ experiment in the presence of 0 mM extracellular Ca^2+^ + 2 mM EGTA or the VDCC blocker verapamil (**Supp. Figure 5B and C)**. We observed no intracellular Ca^2+^ increases in conditions of 0 mM extracellular Ca^2+^ + Li^+^. We also found a significant reduction in the number of the cells responding to Li^++^ verapamil in comparison to Li^+^ alone (61.8 ± 20.62, n= 5, and 9.8 ± 1.9, n= 3, respectively), suggesting that clamping the membrane at E_K_ prevents depolarization and the influx of Ca^2+^ **(Figures 3D and E, Supp. Figure 5B and C**). We conclude that, in the absence of Na^+^ influx through NALCN, SLO2.1 channels are less active, thus reducing K^+^ efflux and depolarizing the MSMC membrane, leading to VDCC activation and Ca^2+^ oscillations.

**Figure 3.**
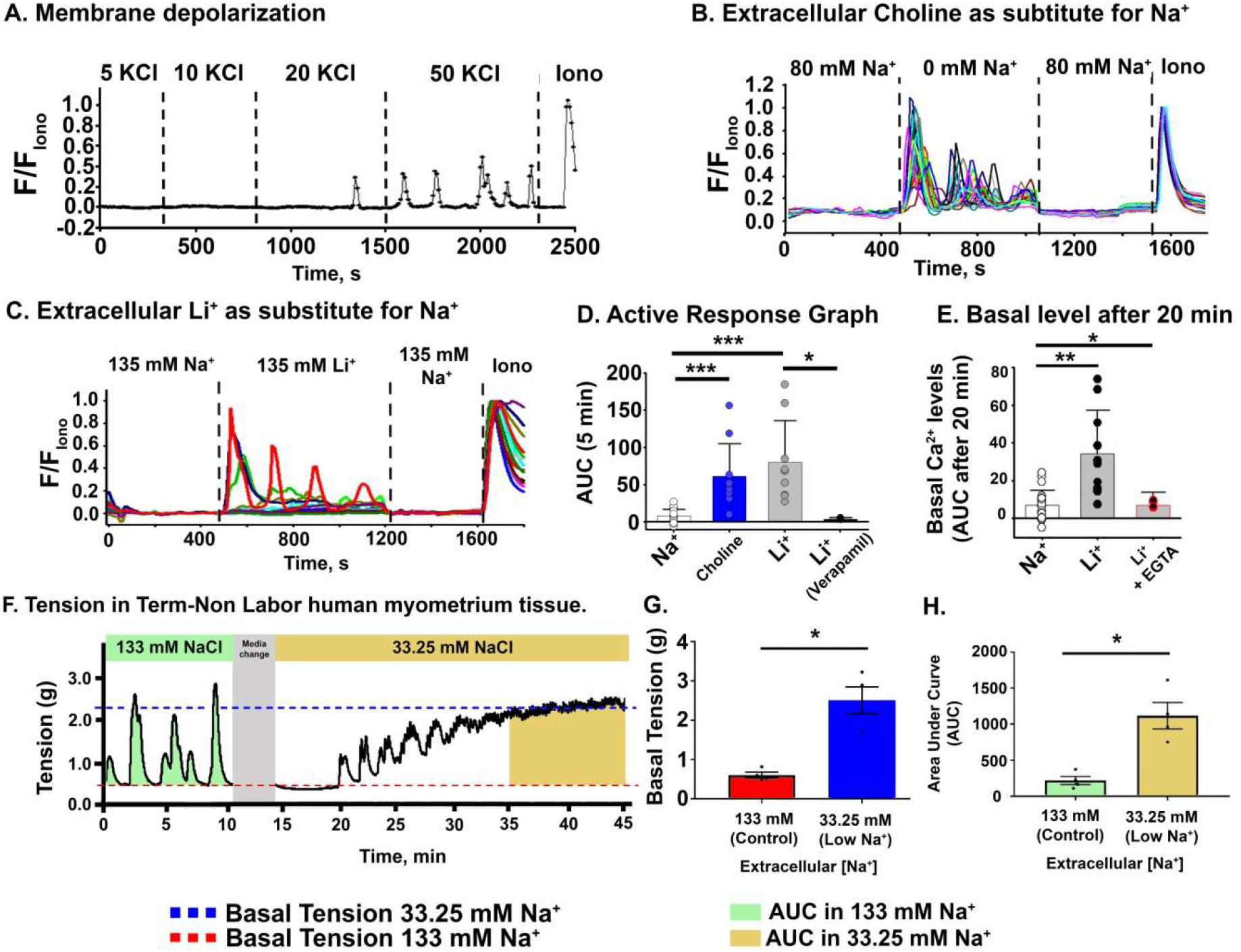
Na^+^ leak regulates intracellular calcium homeostasis and basal tension in human MSMCs and myometrial tissue. **(A, B, C)** Representative fluorescence traces from human MSMCs loaded with 10 μM Fluo-4 AM in the presence of **(A)** 5, 10, 20, and 50 mM external KCl; **(B)** 80 mM Na^+^ or Choline, and **(C)** 135 mM Na^+^ or Li^+^. **(D)** Graphs of the areas under the curve of the first 5 min after changing the solutions in **B, C**, and **Supp. 5C**. In **D**, values are 8.19 ± 8.9 (n=21) for Na^+^, 61.5 ± 43.6 (n=10) for Choline, 80.3 ± 55.8 (n=9) for Li^+^ and 3.3 ± 2.8 (n=3) for Li^+^+ Verapamil (**Supp. 5C**). **(E)** Graphs of the area under the curve (AUC) after 20 min in the indicated solutions. In **E**, values are 7.0 ± 7.7 (n=21) for Na^+^, 34.2 ± 22.9 (n=10) for Li^+^, and 6.94 ± 6.7 (n=4) for Li^+^ + EGTA (**Supp. 5B)**. All data were normalized to the fluorescence in 5 μM ionomycin and 2 mM extracellular Ca^2+^ (Iono), and all data are presented as mean and standard deviation. **(F)** Representative tension-recording trace of myometrial tissue obtained from a woman undergoing elective Cesarean delivery in normal and low extracellular Na^+^. **(G, H)** Quantification of **(G)** basal tension and **(H)** AUC when the tissue was bathed with control or low Na^+^ solutions. Data are presented as mean and standard deviation; n=4 for each. \**P*<0.05, ** *P*<0.01, and *** *P*<0.001 by **(D, E)** unpaired t-test or **(G**,**H)** paired t-test.

In MSMCs, increases in intracellular Ca^2+^ activate MSMC contractile machinery. To determine whether low extracellular Na^+^ would increase uterine muscle basal tension and contractility, we performed tension recording on myometrial strips obtained from women who underwent an elective cesarean delivery. **Figure 3G** shows a representative trace of a myometrial strip bathed in normal Krebs solution containing 133 mM Na^+^. When the extracellular Na^+^ was reduced from 133 mM to 33.25 mM, both the basal (0.61 ± 0.15 g vs. 2.51 ± 0.68 g, *P*=0.0104) and total (219 ± 113 g vs. 1118 ± 366.4 g, *P*=0.0251) tension produced by the myometrial strips significantly increased (**Figures 3H and I**) but rhythmic contractions decreased. This finding supports that Na^+^ influx is needed to activate SLO2.1 in order to maintain the uterus in its resting basal state.

### NALCN and SLO2.1 co-localize in human MSMCs

The above data together suggest that Na^+^ influx through NALCN activates SLO2.1, leading to K^+^ efflux, membrane hyperpolarization, and decreased intracellular Ca^2+^. These findings, combined with the knowledge that SLO2.1 forms functional complexes with the Na^+^ channels that activate it in neurons (Hage & Salkoff, 2012; Takahashi & Yoshino, 2015), led us to propose that NALCN co-localizes with SLO2.1 in human MSMCs. To test this idea, we performed *in* situ proximity ligation assays in both human primary MSMCs and hTERT-HM cells. In this assay, cells are stained with antibodies recognizing the two proteins of interest and DNA-tagged secondary antibodies. If the two proteins are within 40 nm of one another, DNA ligation and amplification occur, which can then be detected as fluorescent punctae in the cells. In both MSMCs and hTERT-HMs, we detected significantly more punctae in cells stained with antibodies specific to NALCN and SLO2.1 than in cells stained with either antibody alone or only secondary antibodies (**Figure 4**). We conclude that NALCN and SLO2.1 are in spatial proximity in human MSMCs.

**Figure 4.**
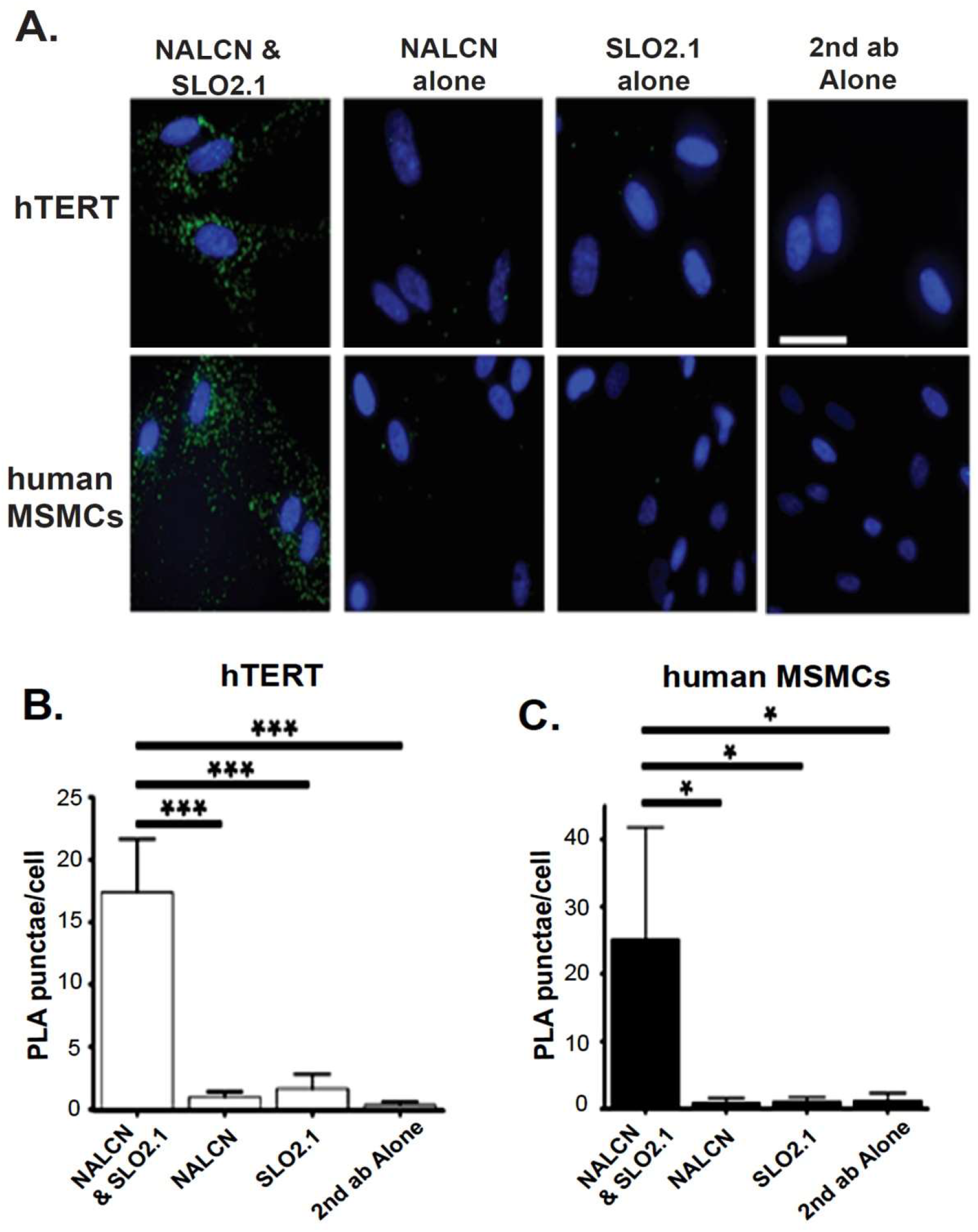
NALCN and SLO2.1 are in proximity in human MSMCs cells. **(A)** Representative proximity ligation assay (PLA) labeling of hTERT-HM and human MSMCs with the indicated single antibodies and antibody combinations. (Scale bar, 10 μm.) **(B, C)** Average number of PLA punctae in **(B)** hTERT-HM cells (n=4) and **(C)** human primary MSMCs (from n=4 patients). Over 300 cells per condition were processed. Data are presented as mean and standard deviation. \**P*<0.05 and \*\*\**P*<0.001 by unpaired t-test.

**Figure 5.**
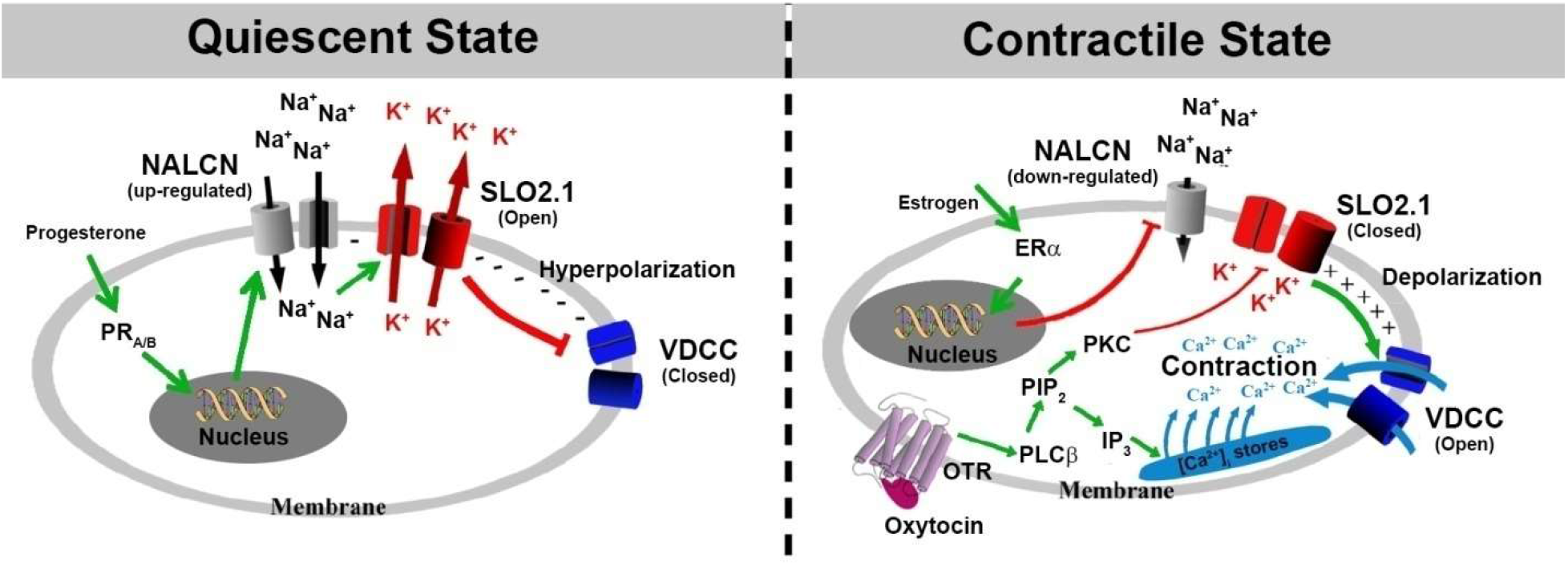
Proposed model by which the NALCN/SLO2.1 complex regulates myometrial excitability. During the quiescent state, progesterone binding to the progesterone receptor (PR_A/B_) increases NALCN expression and activity (Amazu et al., 2020). Sodium current through NALCN activates SLO2.1 channels, increasing K^+^ efflux to maintain the cell in a hyperpolarized state. As a result, voltage-dependent Ca^2+^ channels (VDCCs) are closed, and uterine contractions do not occur. In the contractile state, estrogen acting on ERα inhibits NALCN expression (Amazu et al., 2020), leading to decreased SLO2.1 activity. The reduced K^+^ efflux depolarizes the membrane, leading to VDCC activation, an increase in intracellular Ca^2+^, and uterine contractility. At labor, Oxytocin (OXT) binds to the oxytocin receptor (OTR), leading to activation of phospholipase C (PLC), production of phosphatidylinositol 4,5-bisphosphate (PIP_2_), and production of inositol triphosphate (IP_3_). IP_3_ activates the release of Ca^2+^ from intracellular stores, and PIP_2_ activates protein kinase C (PKC), which inhibits SLO2.1 (Ferreira et al., 2019). This SLO2.1 inhibition further depolarizes the membrane, thus opening more VDCCs, increasing intracellular Ca^2+^, and further activating myosin to cause muscle contraction.

## Discussion

Here, we have presented four lines of evidence that NALCN and SLO2.1 functionally interact to modulate MSMC membrane potential and excitability. First, we report that SLO2.1 activity is activated by Na^+^ influx carried predominately by a NALCN-dependent Na^+^ leak current. Second, this activation of SLO2.1 leads to membrane hyperpolarization, and we show that Na^+^ influx through NALCN is responsible for ∼60% of the SLO2.1-dependent hyperpolarization in human MSMCs. Third, we show that NALCN-mediated regulation of SLO2.1 activity, in turn, regulates Ca^2+^ entry through VDCCs and influences myometrial contractility. Finally, we show that NALCN and SLO2.1 are in spatial proximity in MSMC membranes.

The functional coupling of SLO2.1 K^*+*^ currents to Na^+^ inward currents suggests a highly specialized relationship between SLO2.1 and NALCN channels, resembling the colocalization between calcium channels and the Slo1 calcium-activated potassium channel (BK) (Marrion & Tavalin, 1998). Although the coupling of SLO2.1 to NALCN channels may be analogous, the two orders of magnitude difference in the Na^+^ and Ca^2+^ concentrations required to activate the respective K^+^-channels suggests some differences. One possible mechanism for the functional coupling of SLO2.1 and NALCN currents is their presence in a region of limited diffusion near the plasma membrane. Physiological evidence of such a region (known as fuzzy space) has been reported in cardiac myocytes (Barry, 2006; Semb & Sejersted, 1996). Second, the microheterogeneity of sodium concentrations in internal submembrane space has been directly measured (Wendt-Gallitelli, Voigt, & Isenberg, 1993). Thus, it is possible that a small but constant influx of Na^+^ through NALCN is responsible for increasing [Na^+^] in localized submembrane regions where K_Na_ channels are located. The spatial proximity that we found between SLO2.1 and NALCN channels likely allows NALCN to locally modify the Na^+^ concentration in intracellular microdomains containing SLO2.1 without changing the bulk intracellular Na^+^ concentration (Hage & Salkoff, 2012; P. Li et al., 2019).

We recently published evidence regarding another mechanism by which SLO2.1 is regulated in human MSMCs. In that work, we found that oxytocin acting through a non-canonical pathway inhibits SLO2.1 and induces an increase in intracellular Ca^2+^ by opening VDCC (Ferreira et al., 2019). These results were in line with data published by Arnaudeau *et al*. suggesting that oxytocin has a sustained effect on the VDCC-dependent intracellular Ca^2+^ concentration in MSMCs from pregnant rats (Arnaudeau, Lepretre, & Mironneau, 1994). We also recently demonstrated that NALCN expression and activity are regulated by the hormones progesterone and estrogen (Amazu et al., 2020).

Taken together, our work and the work of others lead us to propose the model shown in **Figure 5**. During the quiescent state, progesterone upregulates the expression of NALCN (Amazu et al., 2020), leading to an increase in the NALCN-dependent Na^+^ leak current. This Na^+^ influx activates SLO2.1 channels locally, leading to K^+^ efflux and membrane hyperpolarization. This hyperpolarization promotes the closed state of VDCCs, limiting Ca^2+^ influx and contractility. During the contractile state, estrogen inhibits the expression of NALCN, reducing Na^+^ influx, and preventing SLO2.1 activation. As a result of decreased SLO2.1-mediated K^+^ efflux, the membrane depolarizes, leading to VDCC activation, Ca^2+^ influx, activation of MSMC contractile machinery, and uterine contractions. Additionally, at labor, oxytocin further induces Ca^2+^ release from intracellular stores and inhibits SLO2.1 activity via signaling through the protein kinase C pathway (Ferreira et al., 2019). Future work will be aimed at further testing this model and defining the mechanisms by which the actions of the hormones progesterone, estrogen, and oxytocin work in concert with the activity of ion channels to regulate MSMC excitability and contractility appropriately during pregnancy.

In addition to SLO2.1, MSMCs express other K^+^ channels including the inward rectifier Kir7.1 and Ca^2+^-activated Slo1.1 channels (Brainard, Korovkina, & England, 2007; Ferreira et al., 2019; Khan et al., 1993; McCloskey et al., 2014). Expression of Kir7.1 channels peaks in mid-pregnancy in mice and Kir7.1 currents contribute to maintaining a hyperpolarized membrane potential and low contractility during the quiescent stage of pregnancy in both mouse and human uterine tissue (McCloskey et al., 2014). This suggests that Kir7.1 could contribute to regulating the transition from the quiescent to the contractile state during pregnancy. However, Ferreira *et al*. showed that Slo1.1 and SLO2.1 together contribute to ∼87% of the K^+^ current in human MSMCs (Ferreira et al., 2019). Further work is thus needed to fully define the activities of the numerous K^+^ channels in MSMCs.

In addition to NALCN, MSMCs from rats and humans also express other channels that conduct Na^+^, such as the TRPC family members TRPC1, 3, and 6 (Babich et al., 2004; Dalrymple et al., 2007; Dalrymple et al., 2002; Ku et al., 2006; Lacampagne, Gannier, Argibay, Garnier, & Le Guennec, 1994; Wang et al., 2020; Yang & Sachs, 1989). Although these channels are inhibited by Gd^3+^, we show here that TRPC1, 3, and 6 jointly contribute to ∼40% of the Na^+^-activated, SLO2.1-dependent hyperpolarization in human MSMCs. The exact regulation and role of TRPCs during pregnancy is under investigation. In one study, TRCP6 mRNA and protein expression are downregulated, whereas TRPC1 expression is unchanged, during pregnancy in rats (Babich et al., 2004). However, others report that TRPCs are sensitive to multiple signals *in vitro*, including mechanical stretch and IL-1β, that are important during labor (Csapo, Erdos, De Mattos, Gramss, & Moscowitz, 1965; Dalrymple et al., 2007; Douglas, Clarke, & Goldspink, 1988). Thus, although TRPCs seem to play some role in regulating MSMC membrane potential, their more important role may be in regulating MSMC activity during labor.

Previous work had shown that depolarizing Na^+^ leak currents could contribute to pacemaker activity in dopaminergic neurons and gastrointestinal cells (Khaliq & Bean, 2010; Kim et al., 2012; Koh, Jun, Kim, & Sanders, 2002). Given that MSMC action potentials are driven by an influx of Ca^2+^ through VDCC that open in response to a slow recurrent depolarization between action potentials (pacemaker current) (Amedee, Mironneau, & Mironneau, 1987; Kuriyama & Suzuki, 1976; Lammers, 2013; Wray et al., 2003), it seemed possible that NALCN could regulate pacemaker activity in MSMCs. However, if NALCN were involved in setting the pacemaker activity, cells lacking NALCN should have a disrupted inter-burst frequency of action potentials. Instead, Reinl *et al*. only observed a significant reduction in the burst duration and no significant effect on the myometrial inter-burst frequency at day 19 of pregnancy in NALCN knockout mice (Reinl et al., 2018). In conclusion, these results suggest that, instead of regulating the pacemaker, NALCN regulates myometrial excitability by modulating the permeability of other ions (particularly K^+^) and thereby regulating MSMC membrane potential (Ferreira et al., 2019; Reinl et al., 2015; Reinl et al., 2018). Additional work is needed to establish the contribution of the functional complex between NALCN and SLO2.1 channels on myometrial excitability at different stages of pregnancy.

## Methods and Materials

### Ethical Approval and Acquisition of Human Samples

This study was approved by the Washington University in St. Louis Institutional Review Board (approval no. 201108143) and conformed to the Declaration of Helsinki except for registration in a database. We obtained signed written consent from each patient. Human tissue samples (0.1-1.0 cm^2^) from the lower uterine segment were obtained from non-laboring women at term (37 weeks of gestation) during elective Cesarean section under spinal anesthesia. Samples were stored at 4°C in phosphate-buffered saline (PBS) and processed for MSMC isolation within 60 min of acquisition.

### Cell Culture

Primary human MSMCs were isolated and cultured as previously described (Y. Li, Lorca, Ma, Rhodes, & England, 2014). Briefly, tissue was treated with PBS containing 50μg/mL gentamicin and 5μg/ml fungizone, then cut into 2 to 3 mm pieces and cultured in DMEM:F12 medium with 5% Fetal Bovine Serum (FBS), 0.2% fibroblast growth factor-β, 0.1% epidermal growth factor, 0.05% insulin, 0.05% gentamicin, and 0.05% fungizone. Colonies were amplified to form primary cell cultures. Primary MSMCs and human telomerase reverse transcriptase-immortalized myometrial cells (hTERT-HM) were incubated at 37°C and 5% CO_2_ in phenol–red-free DMEM: F12 medium with 10% FBS, 100 units/ml penicillin, and 100 µg/ml streptomycin or 25 μg/mL gentamicin (Sigma, St. Louis, MO) (Condon et al., 2002). Primary MSMCs were used within two passages, and hTERT-HM cells were used in passages lower than 15.

### Electrophysiology

Cells were starved in serum-free DMEM:F12 for at least 2 hours before experiments. For all experiments, pipettes were pulled from borosilicate glass from Warner Instruments. For whole-cell recording, pipettes with a resistance of 0.8 to 1.8 megaohms and symmetrical K^+^ were used. External solution was (in mM): 160 KCl, 80 NaCl, 2 MgCl_2_, 10 HEPES, and 5 TEA, pH adjusted to 7.4 with NaOH. For the 0 mM Na^+^ solution, Na^+^ was replaced with 80 mM CholineCl, and pH was adjusted with KOH; the concentration of external K^+^ varied from 4.5 to 5.5 mM. The pipettes were filled with (in mM): 160 KCl, 80 cholineCl or 80 NaCl, 10 HEPES, 0.6 free Mg^2+^, and either 0 or 100 nM free Ca^2+^ solutions with 1 mM EGTA. Variations in the solutions are indicated in the figures. During electrophysiological experiments, the cells and the intracellular side of the membrane were perfused continuously. NALCN inhibitor used for experiments in Figure 1, CP96345 (CP) was obtained from TOCRIS Bioscience, Bristol, UK, Cat # 2893. Traces were acquired with an Axopatch 200B (Molecular Devices), digitized at 10 kHz for whole-cell or macro-patch recordings or at 100 kHz for single-channel recordings. Recordings were filtered at 2 kHz, and pClamp 10.6 (Molecular Devices) and SigmaPlot 12 (Jandel Scientific) were used to analyze the data.

### Determination of Membrane Potential by Flow Cytometry

hTERT-HM cells were centrifuged at 1000 rpm for 5 min. Cells were resuspended in modified Ringers solution containing (in mM): 80 Choline Cl, 10 HEPES, 5 Glucose, 5 KCl, and 2 CaCl_2_; pH 7.4). Before recording, 0.02 mg/mL Hoechst and 150 nM DiSC_3_(5) were added to 500 µL of cell suspension, and data were recorded as individual cellular events on a FACSCanto II TM cytometer (BD Biosciences, Franklin Lakes, NJ). Side scatter area (SSC-A) and forward scatter area (FSC-A) fluorescence data were collected from 100,000 events per recording. Threshold levels for FSC-A and SSC-A were set to exclude signals from cellular debris (Supp. Figure 1A). Doublets, aggregates, and cell debris were excluded from the analysis based on a dual parameter dot plot in which pulse signal (signal high; SSC-H; y-axis) versus signal area (SSC-A; x-axis) was displayed (Supp. Figure 1B). Living Hoechst-negative cells were selected by using the filter Pacific Blue; (450/50), and DiSC3(5)-positive cells were detected with the filter for allophycocyanine APC; (660/20) (Supp. Figure 1C). To measure the effect of Na^+^ on membrane potential, 80 mM NaCl, Choline chloride, or Lithium (Li^+^) was added to the 500 μL suspensions. In some cases, as indicated in the figures, 10 μM gadolinium (Gd^3+^), a concentration known to inhibit NALCN current (Lu et al., 2007), 500 nM GsMTx4 (a peptide inhibitor of TRPC1 and TRPC6 isolated from Grammostolaspatulata spider venom), and 1μM Pyr3 (pyrazole; blocks TRPC3 inhibitor) were added to the cell suspension (Bowman, Gottlieb, Suchyna, Murphy, & Sachs, 2007; Kiyonaka et al., 2009; Lu et al., 2007). Valinomycin hyperpolarizes the cell according to the equilibrium potential for K^+^. Normalization was performed according to (F_Ref_-F_Ion_)/(F_Valino_-F_Ion_), being F_Ref_ median of the fluorescence of the population in the basal condition, F_Ion_ after the addition of the cation, and F_Valino_ after addition of Valinomycin. Normalization was performed by adding 1 µM valinomycin (Sigma, St. Louis, MO). FlowJo 10.6.1 software was used to analyze data, reported as median values.

### Calcium Imaging

Cells were grown on glass coverslips with the media described above for MSMCs. Cells were pre-incubated with 2 μM Fluo-4 AM and 0.05-0.1% Pluronic Acid F-127 in Opti-Mem for 60-90 min. To allow the dye to equilibrate in the cells, the cells were removed from the loading solutions and placed in Ringer solution for 10 to 20 minutes. The various solutions were applied with a perfusion system with an estimated exchange time of 1.5 s. Recordings started 2-5 min before the addition of the first test solution. Ionomycin (5 μM) was added at the end of the recordings as a control stimulus. Calcium (Ca^2+^) signals were recorded with a Leica AF 6000LX system with a Leica DMi8000 inverted microscope and an Andor-Zyla-VCS04494 camera. A halogen lamp was used with a 488 +/-20 nm excitation filter and a 530 +/-20 nm emission filter. A 40X (HC PL FluoTar L 40X/0.70 Dry) or a 20X (N-Plan L 20X/0.35 Dry) air objective were used. Leica LasX 2.0.014332 software was used to collect data and control the system. Acquisition parameters were: 120 ms exposure time, 2×2 binning, 512 x 512 pixels resolution, and a voxel size of 1.3 μm for the 20X objective. Whole images were collected every 10 seconds. LAS X, ImageJ, Clampfit 10 (Molecular Devices), and SigmaPlot 12 were used to analyze data. Changes in intracellular Ca^2+^ concentration are presented as (F/F_Iono_) after background subtraction. All imaging experiments were done at room temperature. Cells were counted as responsive if they had changes in fluorescence of at least 5-10% of ionomycin responses.

### Isometric Tension Recording

Human myometrial tissues (0.1-1.0 cm^2^) from four non-laboring women at term were isolated and cut into strips (10 x 2 mm) and immediately placed in 4° Krebs solution containing (in mM): 133 NaCl, 4.7 KCl, 1.2 MgSO_4_, 1.2 KH_2_PO_4_, 10 TES, 1.2 CaCl_2_, and 11.1 glucose, pH 7.4. Strips were mounted to a force transducer in organ baths filled with oxygenated (95% O_2_, 5% CO_2_) Krebs solution at 35.7°C, and tension was recorded with a muscle strip myographdata acquisition system (DMT, Ann Arbor, MI). Basal tension (2 g) was applied to the tissue strips, and strips were equilibrated for 1 hr until spontaneous myometrial contractility appeared. When four stable regular contractile waveforms were observed, Krebs solution was changed to modified Krebs solution containing (in mM): 33.25 NaCl, 99.75 CholineCl, 4.7 KCl, 1.2 MgSO_4_, 1.2 KH_2_PO_4_, 10 TES, 1.2 CaCl_2_, and 11.1 glucose, pH 7.4. Tension was recorded for 30 min. Traces obtained in the last 10 minutes before and after adding modified Krebs solution were compared by using LabChart 8 (ADInstruments, Colorado Springs, CO). Basal tension and area under the curve (AUC) of phasic contractions of the myometrial strips were calculated in both solutions.

### *In Situ* Proximity Ligation Assay

The hTERT-HM cells were cultured in 8-well chambered slides (LabTek/Sigma, St. Louis, MO), serum-deprived in 0.5% FBS for 24 h, washed in ice-cold 1X PBS, and then fixed in 4% (wt/vol) paraformaldehyde (PFA) in PBS for 20 min at room temperature with gentle rocking. After 4 x 5-min washes in 1X PBS, cells were permeabilized with 0.1% NP-40 for 5 min at room temperature, washed twice with PBS, and washed once in 100 mM Glycine in PBS to quench remaining PFA. The slides were rinsed in milliQ water to remove residual salts. Duolink*in situ* proximity ligation assay (Sigma, St. Louis, MO) labeling was performed with the following antibodies: NALCN (mouse monoclonal, 1:100, StressMarq) and SLO2.1 (rabbit polyclonal, 1:200, Alomone). The manufacturer’s protocol was followed completely except that cells were stained with NucBlue Fixed Cell Stain ReadyProbes (Invitrogen, Carlsbad, CA) for 5 min at room temperature before the final wash in wash buffer B. The slides were dried at room temperature in the dark, mounted in Vectashield (Vector Laboratories, Burlingame, CA), and stored in the dark at -20 °C until analysis. Images were collected with a fluorescent Leica AF 6000LX system (Buffalo Grove, IL, USA) with a Leica DMi8000 inverted microscope and an Andor-Zyla-VCS04494 camera with excitation wavelengths of 488 nm for PLA signals and 340 nm for NucBlue. A 63X objective (HC PL FluoTar L 63X/0.70 Dry) was used to obtain the images. Leica LasX software was used to control the system and collect data. Acquisition parameters were: 2 and 1.5 seconds of exposure time for 488 and 340 nm, respectively, no binning, 2048 x 2048 pixels resolution, and a voxel size of 0.103 μm. Whole images were collected every 10 seconds. LAS X, ImageJ software (National Institutes of Health, Bethesda, Maryland, USA), and SigmaPlot 12 (Systat Software Inc., Chicago, IL, USA) were used to analyze images. Data are presented as number of punctae (counted after background subtraction) per cell. All imaging experiments were done at room temperature.

### Statistical Analyses

Sigmaplot, version 12.0 (Systat Software Inc.) was used for all statistical analyses. An unpaired Student’s t-test was used to compare independent samples, and a paired t-test was used to compare data in case-control studies performed in the same individuals. Data are expressed as the mean ± SD. *P-value* < 0.05 was considered statistically significant.

## Acknowledgments

We thank Dr. Deborah Frank for the critical review of the manuscript. We also thank the Clinical Research Nurses in the Department of Obstetrics and Gynecology at Barnes Jewish Hospital for consenting patients and acquiring human myometrial biopsies. We thank the Washington University Flow Cytometry & Fluorescence-Activated Cell Sorting Cores for use of their FACSCanto II TM cytometer and guidance.

## Author Contributions

J.J.F. and C.A. designed, performed, and analyzed the experiments. L.C.P.M. assisted with the flow cytometry experiments. X.M performed experiments measuring the inhibition of Na^+^ leak currents by CP96345. J.J.F., C.A., S.K.E., and C.M.S. analyzed data and interpreted the results. J.J.F. and C.A. prepared figures. J.J.F., C.A., S.K.E., and C.M.S drafted the manuscript. J.J.F., C.A., L.C.P.M., X.M, S.K.E., and C.M.S. edited, revised, and approved the final version of the manuscript.

## References

Amazu, C., Ma, X., Henkes, C., Ferreira, J. J., Santi, C. M., & England, S. K. (2020). Progesterone and estrogen regulate NALCN expression in human myometrial smooth muscle cells. [Research Support, N.I.H., Extramural Research Support, Non-U.S. Gov’t]. Am J Physiol Endocrinol Metab, 318(4), E441–E452. doi: 10.1152/ajpendo.00320.2019

Amedee, T., Mironneau, C., & Mironneau, J. (1987). The calcium channel current of pregnant rat single myometrial cells in short-term primary culture. [Research Support, Non-U.S. Gov’t]. J Physiol, 392, 253–272. doi: 10.1113/jphysiol.1987.sp016779

Arnaudeau, S., Lepretre, N., & Mironneau, J. (1994). Oxytocin mobilizes calcium from a unique heparin-sensitive and thapsigargin-sensitive store in single myometrial cells from pregnant rats. [Research Support, Non-U.S. Gov’t]. Pflugers Arch, 428(1), 51–59. doi: 10.1007/BF00374751

Babich, L. G., Ku, C. Y., Young, H. W., Huang, H., Blackburn, M. R., & Sanborn, B. M. (2004). Expression of capacitative calcium TrpC proteins in rat myometrium during pregnancy. [Research Support, U.S. Gov’t, P.H.S.]. Biol Reprod, 70(4), 919–924. doi: 10.1095/biolreprod.103.023325

Barry, W. H. (2006). Na”Fuzzy space”: does it exist, and is it important in ischemic injury? [Comment Editorial Research Support, Non-U.S. Gov’t Review]. J Cardiovasc Electrophysiol, 17 Suppl 1, S43–S46. doi: 10.1111/j.1540-8167.2005.00396.x

Bowman, C. L., Gottlieb, P. A., Suchyna, T. M., Murphy, Y. K., & Sachs, F. (2007). Mechanosensitive ion channels and the peptide inhibitor GsMTx-4: history, properties, mechanisms and pharmacology. [Review]. Toxicon, 49(2), 249–270. doi: 10.1016/j.toxicon.2006.09.030

Brainard, A. M., Korovkina, V. P., & England, S. K. (2007). Potassium channels and uterine function. [Research Support, N.I.H., Extramural Research Support, Non-U.S. Gov’t Review]. Semin Cell Dev Biol, 18(3), 332–339. doi: 10.1016/j.semcdb.2007.05.008

Budelli, G., Hage, T. A., Wei, A., Rojas, P., Jong, Y. J., O’Malley, K., & Salkoff, L. (2009). Na+-activated K+ channels express a large delayed outward current in neurons during normal physiology. [Research Support, N.I.H., Extramural]. Nat Neurosci, 12(6), 745–750. doi: 10.1038/nn.2313

Casteels, R., & Kuriyama, H. (1965). Membrane Potential and Ionic Content in Pregnant and Non-Pregnant Rat Myometrium. J Physiol, 177, 263–287. doi: 10.1113/jphysiol.1965.sp007591

Condon, J., Yin, S., Mayhew, B., Word, R. A., Wright, W. E., Shay, J. W., & Rainey, W. E. (2002). Telomerase immortalization of human myometrial cells. [Research Support, U.S. Gov’t, P.H.S.]. Biol Reprod, 67(2), 506–514. doi: 10.1095/biolreprod67.2.506

Csapo, A., Erdos, T., De Mattos, C. R., Gramss, E., & Moscowitz, C. (1965). Stretch-induced uterine growth, protein synthesis and function. Nature, 207(5004), 1378–1379. doi: 10.1038/2071378a0

Dalrymple, A., Mahn, K., Poston, L., Songu-Mize, E., & Tribe, R. M. (2007). Mechanical stretch regulates TRPC expression and calcium entry in human myometrial smooth muscle cells. [Research Support, Non-U.S. Gov’t]. Mol Hum Reprod, 13(3), 171–179. doi: 10.1093/molehr/gal110

Dalrymple, A., Slater, D. M., Beech, D., Poston, L., & Tribe, R. M. (2002). Molecular identification and localization of Trp homologues, putative calcium channels, in pregnant human uterus. [Research Support, Non-U.S. Gov’t]. Mol Hum Reprod, 8(10), 946–951. doi: 10.1093/molehr/8.10.946

Douglas, A. J., Clarke, E. W., & Goldspink, D. F. (1988). Influence of mechanical stretch on growth and protein turnover of rat uterus. Am J Physiol, 254(5 Pt 1), E543-548. doi: 10.1152/ajpendo.1988.254.5.E543

Dryer, S. E. (2003). Molecular identification of the Na+-activated K+ channel. [Comment]. Neuron, 37(5), 727–728. doi: 10.1016/s0896-6273(03)00119-3

Ferreira, J. J., Butler, A., Stewart, R., Gonzalez-Cota, A. L., Lybaert, P., Amazu, C., … Santi, C. M. (2019). Oxytocin can regulate myometrial smooth muscle excitability by inhibiting the Na(+) -activated K(+) channel, Slo2.1. [Research Support, N.I.H., Extramural Research Support, Non-U.S. Gov’t]. J Physiol, 597(1), 137–149. doi: 10.1113/JP276806

Hage, T. A., & Salkoff, L. (2012). Sodium-activated potassium channels are functionally coupled to persistent sodium currents. [Research Support, N.I.H., Extramural]. J Neurosci, 32(8), 2714–2721. doi: 10.1523/JNEUROSCI.5088-11.2012

Hahn, S., Kim, S. W., Um, K. B., Kim, H. J., & Park, M. K. (2020). N-benzhydryl quinuclidine compounds are a potent and Src kinase-independent inhibitor of NALCN channels. Br J Pharmacol, 177(16), 3795–3810. doi: 10.1111/bph.15104

Kameyama, M., Kakei, M., Sato, R., Shibasaki, T., Matsuda, H., & Irisawa, H. (1984). Intracellular Na+ activates a K+ channel in mammalian cardiac cells. [Research Support, Non-U.S. Gov’t]. Nature, 309(5966), 354–356. doi: 10.1038/309354a0

Khaliq, Z. M., & Bean, B. P. (2010). Pacemaking in dopaminergic ventral tegmental area neurons: depolarizing drive from background and voltage-dependent sodium conductances. [Research Support, N.I.H., Extramural Research Support, Non-U.S. Gov’t]. J Neurosci, 30(21), 7401–7413. doi: 10.1523/JNEUROSCI.0143-10.2010

Khan, R. N., Smith, S. K., Morrison, J. J., & Ashford, M. L. (1993). Properties of large-conductance K+ channels in human myometrium during pregnancy and labour. [Research Support, Non-U.S. Gov’t]. Proc Biol Sci, 251(1330), 9–15. doi: 10.1098/rspb.1993.0002

Kim, B. J., Chang, I. Y., Choi, S., Jun, J. Y., Jeon, J. H., Xu, W. X., … So, I. (2012). Involvement of Na(+)-leak channel in substance P-induced depolarization of pacemaking activity in interstitial cells of Cajal. [Research Support, N.I.H., Extramural Research Support, Non-U.S. Gov’t]. Cell Physiol Biochem, 29(3-4), 501–510. doi: 10.1159/000338504

Kiyonaka, S., Kato, K., Nishida, M., Mio, K., Numaga, T., Sawaguchi, Y., … Mori, Y. (2009). Selective and direct inhibition of TRPC3 channels underlies biological activities of a pyrazole compound. [Research Support, Non-U.S. Gov’t]. Proc Natl Acad Sci U S A, 106(13), 5400–5405. doi: 10.1073/pnas.0808793106

Koh, S. D., Jun, J. Y., Kim, T. W., & Sanders, K. M. (2002). A Ca(2+)-inhibited non-selective cation conductance contributes to pacemaker currents in mouse interstitial cell of Cajal. [Research Support, Non-U.S. Gov’t Research Support, U.S. Gov’t, P.H.S.]. J Physiol, 540(Pt 3), 803–814. doi: 10.1113/jphysiol.2001.014639

Ku, C. Y., Babich, L., Word, R. A., Zhong, M., Ulloa, A., Monga, M., & Sanborn, B. M. (2006). Expression of transient receptor channel proteins in human fundal myometrium in pregnancy. [Research Support, N.I.H., Extramural]. J Soc Gynecol Investig, 13(3), 217–225. doi: 10.1016/j.jsgi.2005.12.007

Kuriyama, H., & Suzuki, H. (1976). Changes in electrical properties of rat myometrium during gestation and following hormonal treatments. J Physiol, 260(2), 315–333. doi: 10.1113/jphysiol.1976.sp011517

Lacampagne, A., Gannier, F., Argibay, J., Garnier, D., & Le Guennec, J. Y. (1994). The stretch-activated ion channel blocker gadolinium also blocks L-type calcium channels in isolated ventricular myocytes of the guinea-pig. [Research Support, Non-U.S. Gov’t]. Biochim Biophys Acta, 1191(1), 205–208. doi: 10.1016/0005-2736(94)90250-x

Lammers, W. J. (2013). The electrical activities of the uterus during pregnancy. [Research Support, Non-U.S. Gov’t Review]. Reprod Sci, 20(2), 182–189. doi: 10.1177/1933719112446082

Li, P., Halabi, C. M., Stewart, R., Butler, A., Brown, B., Xia, X., … Salkoff, L. (2019). Sodium-activated potassium channels moderate excitability in vascular smooth muscle. [Research Support, N.I.H., Extramural]. J Physiol, 597(20), 5093–5108. doi: 10.1113/JP278279

Li, Y., Lorca, R. A., Ma, X., Rhodes, A., & England, S. K. (2014). BK channels regulate myometrial contraction by modulating nuclear translocation of NF-kappaB. [Research Support, N.I.H., Extramural Research Support, Non-U.S. Gov’t]. Endocrinology, 155(8), 3112–3122. doi: 10.1210/en.2014-1152

Lu, B., Su, Y., Das, S., Liu, J., Xia, J., & Ren, D. (2007). The neuronal channel NALCN contributes resting sodium permeability and is required for normal respiratory rhythm. [Research Support, Non-U.S. Gov’t]. Cell, 129(2), 371–383. doi: 10.1016/j.cell.2007.02.041

Lynn, S., Morgan, J. M., Gillespie, J. I., & Greenwell, J. R. (1993). A novel ryanodine sensitive calcium release mechanism in cultured human myometrial smooth-muscle cells. [Research Support, Non-U.S. Gov’t]. FEBS Lett, 330(2), 227–230. doi: 10.1016/0014-5793(93)80279-4

Marrion, N. V., & Tavalin, S. J. (1998). Selective activation of Ca2+-activated K+ channels by co-localized Ca2+ channels in hippocampal neurons. [Research Support, U.S. Gov’t, P.H.S.]. Nature, 395(6705), 900–905. doi: 10.1038/27674

McCloskey, C., Rada, C., Bailey, E., McCavera, S., van den Berg, H. A., Atia, J., … Blanks, A. M. (2014). The inwardly rectifying K+ channel KIR7.1 controls uterine excitability throughout pregnancy. [Research Support, N.I.H., Extramural Research Support, Non-U.S. Gov’t]. EMBO Mol Med, 6(9), 1161–1174. doi: 10.15252/emmm.201403944

Molina, L. C. P., Gunderson, S., Riley, J., Lybaert, P., Borrego-Alvarez, A., Jungheim, E. S., & Santi, C. M. (2019). Membrane Potential Determined by Flow Cytometry Predicts Fertilizing Ability of Human Sperm. Front Cell Dev Biol, 7, 387. doi: 10.3389/fcell.2019.00387

Morgan, J. M., Lynn, S., Gillespie, J. I., & Greenwell, J. R. (1993). Measurements of Intracellular Ca2+ in Cultured Human Myometrial Smooth-Muscle Cells Bathed in Low Na+ Solutions. Experimental Physiology, 78(5), 711–714. doi: DOI 10.1113/expphysiol.1993.sp003719

Parkington, H. C., Tonta, M. A., Brennecke, S. P., & Coleman, H. A. (1999). Contractile activity, membrane potential, and cytoplasmic calcium in human uterine smooth muscle in the third trimester of pregnancy and during labor. [Research Support, Non-U.S. Gov’t]. Am J Obstet Gynecol, 181(6), 1445–1451. doi: 10.1016/s0002-9378(99)70390-x

Plasek, J., & Hrouda, V. (1991). Assessment of membrane potential changes using the carbocyanine dye, diS-C3-(5): synchronous excitation spectroscopy studies. Eur Biophys J, 19(4), 183–188. doi: 10.1007/BF00196344

Reinl, E. L., Cabeza, R., Gregory, I. A., Cahill, A. G., & England, S. K. (2015). Sodium leak channel, non-selective contributes to the leak current in human myometrial smooth muscle cells from pregnant women. [Research Support, N.I.H., Extramural Research Support, Non-U.S. Gov’t]. Mol Hum Reprod, 21(10), 816–824. doi: 10.1093/molehr/gav038

Reinl, E. L., Zhao, P., Wu, W., Ma, X., Amazu, C., Bok, R., … England, S. K. (2018). Na+-Leak Channel, Non-Selective (NALCN) Regulates Myometrial Excitability and Facilitates Successful Parturition. Cell Physiol Biochem, 48(2), 503–515. doi: 10.1159/000491805

Santi, C. M., Martinez-Lopez, P., de la Vega-Beltran, J. L., Butler, A., Alisio, A., Darszon, A., & Salkoff, L. (2010). The SLO3 sperm-specific potassium channel plays a vital role in male fertility. [Research Support, N.I.H., Extramural Research Support, Non-U.S. Gov’t]. FEBS Lett, 584(5), 1041–1046. doi: 10.1016/j.febslet.2010.02.005

Semb, S. O., & Sejersted, O. M. (1996). Fuzzy space and control of Na+, K(+)-pump rate in heart and skeletal muscle. [Research Support, Non-U.S. Gov’t Review]. Acta Physiol Scand, 156(3), 213–225. doi: 10.1046/j.1365-201X.1996.211000.x

Smith, C. O., Wang, Y. T., Nadtochiy, S. M., Miller, J. H., Jonas, E. A., Dirksen, R. T., … Brookes, P. S. (2018). Cardiac metabolic effects of KNa1.2 channel deletion and evidence for its mitochondrial localization. FASEB J, fj201800139R. doi: 10.1096/fj.201800139R

Takahashi, I., & Yoshino, M. (2015). Functional coupling between sodium-activated potassium channels and voltage-dependent persistent sodium currents in cricket Kenyon cells. [Research Support, Non-U.S. Gov’t]. J Neurophysiol, 114(4), 2450–2459. doi: 10.1152/jn.00087.2015

Wang, H., Cheng, X., Tian, J., Xiao, Y., Tian, T., Xu, F., … Zhu, M. X. (2020). TRPC channels: Structure, function, regulation and recent advances in small molecular probes. [Review Research Support, Non-U.S. Gov’t Research Support, N.I.H., Extramural]. Pharmacol Ther, 209, 107497. doi: 10.1016/j.pharmthera.2020.107497

Wendt-Gallitelli, M. F., Voigt, T., & Isenberg, G. (1993). Microheterogeneity of subsarcolemmal sodium gradients. Electron probe microanalysis in guinea-pig ventricular myocytes. J Physiol, 472, 33–44. doi: 10.1113/jphysiol.1993.sp019934

Wray, S., Jones, K., Kupittayanant, S., Li, Y., Matthew, A., Monir-Bishty, E., … Shmygol, A. V. (2003). Calcium signaling and uterine contractility. [Research Support, Non-U.S. Gov’t Review]. J Soc Gynecol Investig, 10(5), 252–264. doi: 10.1016/s1071-5576(03)00089-3

Yang, X. C., & Sachs, F. (1989). Block of stretch-activated ion channels in Xenopus oocytes by gadolinium and calcium ions. [Comparative Study Research Support, Non-U.S. Gov’t Research Support, U.S. Gov’t, Non-P.H.S. Research Support, U.S. Gov’t, P.H.S.]. Science, 243(4894 Pt 1), 1068-1071. doi: 10.1126/science.2466333

Yuan, A., Santi, C. M., Wei, A., Wang, Z. W., Pollak, K., Nonet, M., … Salkoff, L. (2003). The sodium-activated potassium channel is encoded by a member of the Slo gene family. [Research Support, Non-U.S. Gov’t Research Support, U.S. Gov’t, P.H.S.]. Neuron, 37(5), 765–773. doi: 10.1016/s0896-6273(03)00096-5

