## Supplementary Figures for "A novel sodium signaling complex regulates uterine activity"

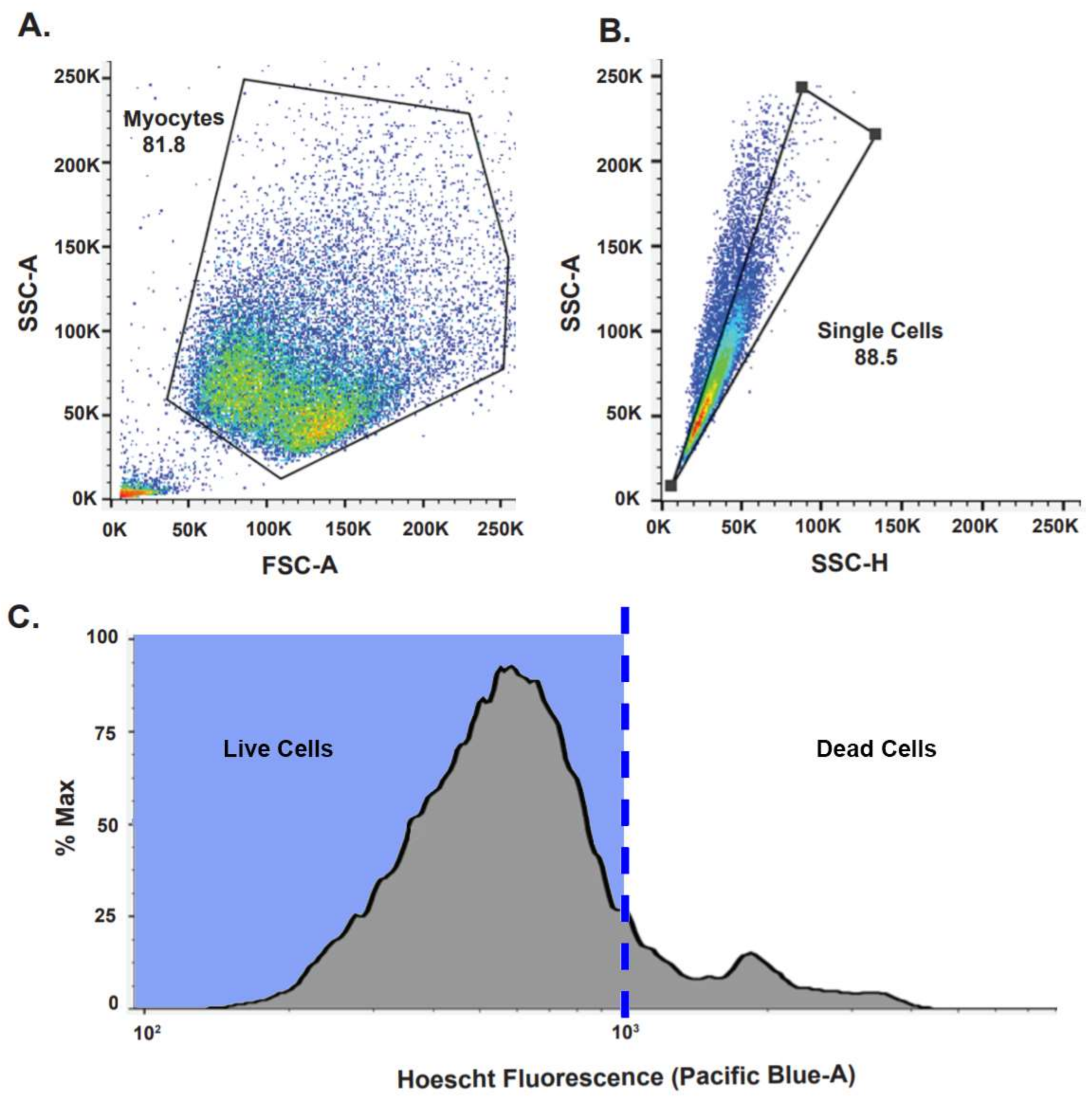


**Supplementary Figure 1: Optimized flow cytometry parameters. (A)** SSC-A and FSC-A were used to identify MSMCs primarily by size. **(B)** SSC-A and SSC-H were used to select singlets and exclude doublets or debris. **(C)** Hoechst dye was used to differentiate between live and dead MSMCs.

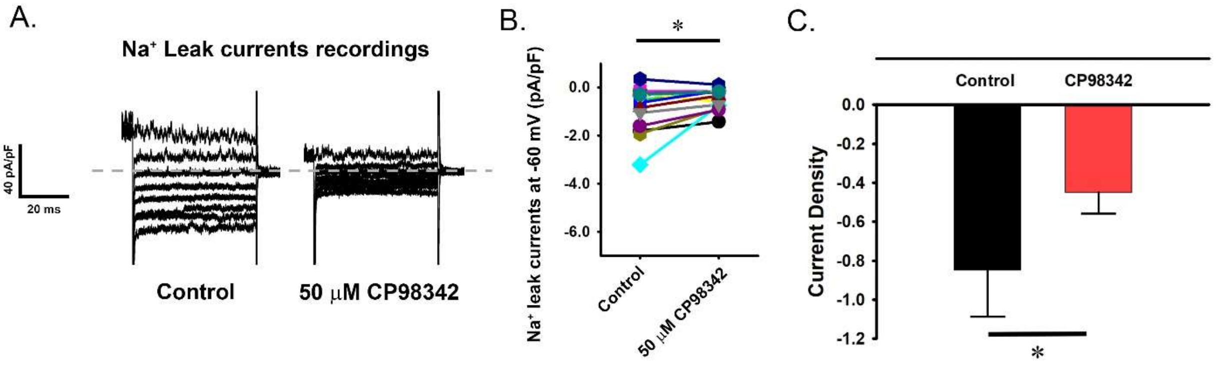


**Supplementary Figure 2. $Na^+$ leak currents inhibited by CP96345 in hTERT Cells. (A)** Representative traces of leak current evoked from hTERT cells. Cells were held at 0 mV and currents were elicited by using a voltage step protocol with 20 mV increments from -100 mV to +20 mV, before and after treatment with 50 μM CP96345 (CP). Capacitive currents were removed from these traces. **(B)** $Na^+$ leak currents at -60 mV obtained before (control) and after treatment with 50 μM CP. Symbols represent individual cells and connecting lines link values in the same cell sample before and after treatment. **(C)** Current density analysis before and after treatment with 50 μM CP. Data are presented as mean values and SD. Here experiments cells were serum-starved in 0.5% FBS-containing media for 20–24 hours before experiments were performed. Pipettes were filled with a solution containing 125 mM Cs-Aspartate, 20 mM tetraethylammonium (TEA)-Cl, 5 mM Mg-ATP, 5 mM EGTA, 100 nM free $Ca^{2+}$, and 10 mM HEPES, pH 7.2. Currents were measured in an extracellular solution containing: 125 mM NaCl, 20 mM TEA-Cl, 0.1 mM $MgCl_2$, 5 mM HEPES, 11 mM glucose, 1 mM $CaCl_2$, and 5 mM nifedipine, pH 7.4.

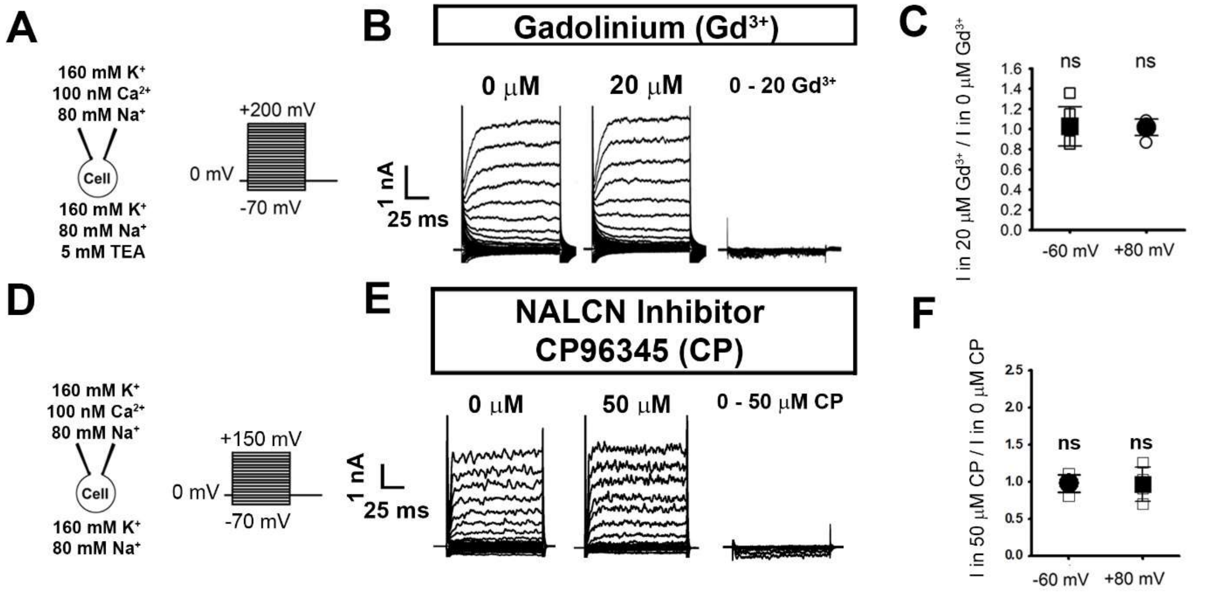


**Supplementary Figure 3. Gadolinium and CP96345 do not directly inhibit SLO2.1 currents. (A)** Schematic of whole-cell recording set-up and voltage steps applied. **(B)** Representative whole-cell currents ($V_h$ = 0 mV, with step pulses from −80 to +150 mV) from human MSMC recorded in 0 or 20 μM $Gd^{3+}$. **(C)** Graph of currents in the presence of 20 μM $Gd^{3+}$ at -60 mV and +80 mV, normalized to currents in 0 μM $Gd^{3+}$ and presented as mean and standard deviation. Values are 1.03 ± 0.195 at -60mV (n=6, $P$ = 0.739) and 1.018 ± 0.0814 at +80 mV (n= 6, $P$ = 0.589). **(D)** Schematic of whole-cell recording set-up and voltage steps applied. **(E)** Representative whole-cell currents ($V_h$ = 0 mV, with step pulses from −80 to +150 mV) from human MSMC recorded in 0 or 50 μM CP. **(F)** Graph of currents in the presence of 50 μM CP at -60 mV and +80 mV, normalized to currents in 0 μM CP and presented as mean and standard deviation. Values are 0.976 ± 0.119 at -60mV (n=6) and 0.972 ± 0.237 at +80 mV (n= 6). $P$-values calculated by paired t-test. ns, not significant.

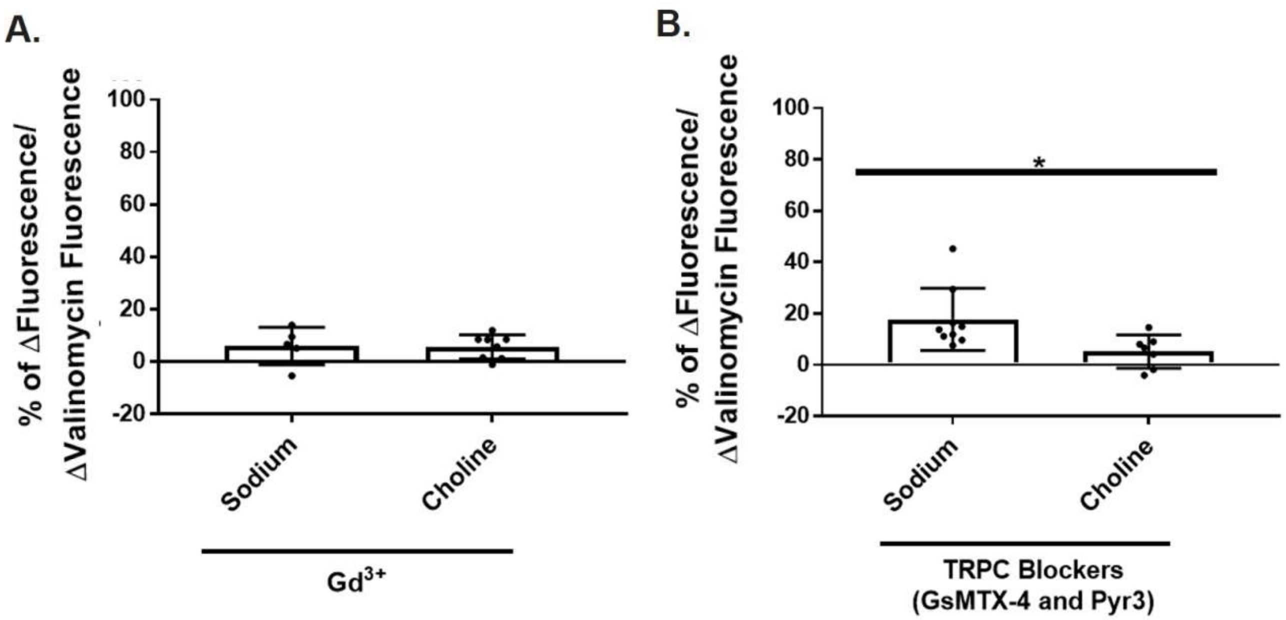


**Supplementary Figure 4: Gadolinium blocks hyperpolarization caused by both extracellular $Na^+$ and Choline, whereas TRPC blockers only partially decrease the hyperpolarization caused by extracellular $Na^+$. (A)** Quantification of shifts in DiSC3(5) fluorescence caused by Sodium (n= 5) and Choline (n=6) in the presence of $Gd^{3+}$. **(B)** Quantification of shifts in DiSC3(5) fluorescence caused by Sodium (n=9) and Choline (n=7) in the presence of GsMTx-4 and Pyr3. In **(A)** and **(B)**, values are normalized to shifts in fluorescence in the presence of valinomycin. Data are presented as mean and standard deviation. *$P<0.05$ by unpaired t-test with Mann-Whitney corrections.

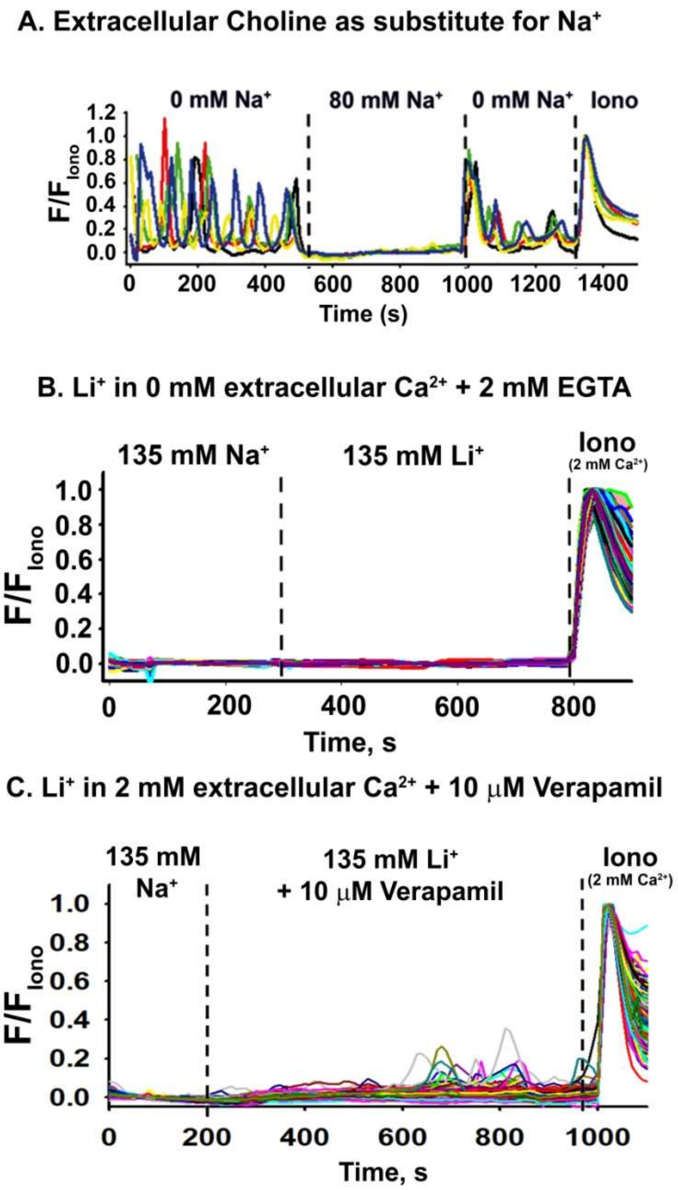


**Supplementary Figure 5: $Na^+$ leak regulates intracellular calcium homeostasis and basal tension in human MSMCs. (A, B, and C)** Representative fluorescence traces from human MSMCs loaded with 10 μM Fluo-4 AM. **(A)** Calcium responses in MSMCs by substituting extracellular Choline with $Na^+$. Reverse experiments of Figure 4B. **(B)** Calcium responses in extracellular 0 mM $Ca^{2+}$ + 2 mM EGTA, when substituting extracellular $Na^+$ with $Li^+$ (Values represented in Figure 3E). **(C)** Calcium responses on cells bathed with 10 μM Verapamil, when substituting extracellular $Na^+$ with $Li^+$ (Values on Figure 3D). All data were normalized to the fluorescence in 5 μM ionomycin and 2 mM extracellular $Ca^{2+}$ (Iono).
